# Genomic markers of drug resistance in *Mycobacterium tuberculosis* populations with minority variants

**DOI:** 10.1101/2023.04.19.537555

**Authors:** Xiaomei Zhang, Connie Lam, Elena Martinez, Eby Sim, Taryn Crighton, Ben J Marais, Vitali Sintchenko

## Abstract

Minority variants of *Mycobacterium tuberculosis* harbouring mutations conferring resistance can become dominant populations during tuberculosis (TB) treatment, leading to treatment failure. Our understanding of drug resistant within-host sub-populations and the frequency of resistance conferring mutations in minority variants remains limited.

*M. tuberculosis* sequences recovered from liquid cultures of culture-confirmed TB cases notified between January 2017 and December 2021 in New South Wales, Australia were examined. Potential drug resistance conferring minority variants were identified using LoFreq, and mixed populations of different *M. tuberculosis* strains (≥100 SNPs apart) were examined using QuantTB.

A total of 1831 routinely sequenced *M. tuberculosis* strains were included in the analysis. Drug resistance conferring minority variants were detected in 3.5% (65/1831) of sequenced cultures; 84.6% (55/65) had majority strains that were drug susceptible and 15.4% (10/65) had majority strains that were drug resistant. Minority variants with high confidence drug resistance conferring mutations were 1.5 times more common when the majority strains were drug resistant. Mixed *M. tuberculosis* strain populations were documented in 10.0% (183/1831) of specimens. Minority variants with high confidence drug resistance conferring mutations were more frequently detected in mixed *M. tuberculosis* strain populations (2.7%, 5/183) than in single strain populations (0.6%, 10/1648; p=0.01).

Drug resistant minority variants require careful monitoring in settings that implement routine *M. tuberculosis* sequencing. The frequency with which drug resistant minority variants are detected is influenced by selective culture methods and culture-independent sequencing should provide a more accurate picture.

## Introduction

The emergence and spread of drug-resistant (DR) *Mycobacterium tuberculosis* strains pose a major global public health challenge (1, 2). The World Health Organization (WHO) estimated that 450 000 new tuberculosis (TB) cases had rifampicin-resistant (RR) or multidrug-resistant (MDR) TB in 2021, although only 166 991 (37.1%)) were formally reported (1). Undetected drug resistant TB (DR-TB) is a major concern, since it compromises patient care and facilitates DR-TB transmission (1).

The presence of drug resistant *M. tuberculosis* strains as minority sub-populations in diagnostic samples has been documented (3, 4). These minority sub-populations may represent either ‘mixed infection’ (simultaneous co-infection with more than one strain) or minority variants of the same strain. The presence of minority variants with drug resistance conferring mutations could promote resistance amplification if patients receive a sub-optimal treatment (5), compromising patient outcomes and facilitating the spread of these strains within the community (6-8).

The detection of drug resistant *M. tuberculosis* can be achieved using either culture-based phenotypic drug-susceptibility testing (pDST) or genotypic drug resistance prediction (gDST), which relies on rapid molecular tests or whole genome sequencing (WGS) to detect mutations associated with drug resistance (1, 9). Culture-based pDST can potentially detect drug resistant sub-populations that are present at more than 1% of the cultured population (8, 10). Several rapid gDST methods have been endorsed by the WHO to detect drug resistance directly from clinical specimens, without initial culture. Both Xpert^®^ MTB/RIF and MTB/RIF Ultra are reported to be able to detect rifampicin-resistance if present at more than 20% of *M. tuberculosis* population in the sample, while GenoTypeMTBDR*plus*v2.0 (LPA-Hain) detects isoniazid or rifampicin resistance if present in more than 5% of bacteria at the site of infection (4, 11, 12). All molecular tests are limited by the fact that they can only assess a small number of pre-determined mutations, which is not inclusive of many mutations associated with drug resistance listed in the WHO Catalogue (13).

WGS studies have demonstrated the presence of drug resistant sub-populations in clinical samples (14, 15), confirming results from earlier molecular strain-typing methods (3, 4). However, WGS analysis of drug-resistance mutations usually focus on the majority population present within the diagnostic specimen (16, 17), but recent validation of genome-wide variant calling tools for the detection of minority resistance-conferring variants have opened new opportunities to identify mixed populations and minority variants in raw sequencing data (17, 18). Specifically, Lofreq allows detection of minority variants harbouring high-confidence drug-resistance conferring mutations that may not be recognised by standard WGS data analysis pipelines (17, 18). In instances with multi-strain infection, QuantTB (19) quantifies the estimated mixture of constituent strains based on a strain specific single-nucleotide polymorphism (SNP) reference database and SNP differences between strains.

This study leveraged our prospective sequencing of all culture-confirmed TB cases and aimed to investigate the incidence of drug-resistance conferring minority variants and mixed strain infections. Accurate detection of minority variants with associated drug-resistance, whether mixed strain or same strain variants, is important for optimal clinical care and public health surveillance.

## Methods

Genomic sequences of *M. tuberculosis* cultures from TB cases notified between 1 January 2017 and 31 December 2021 in New South Wales, Australia were examined, together with their respective pDST results (Supplementary Table S1).

### Detection of minority variants with drug resistance

The reference genome used for the genomic analysis was *M. tuberculosis* strain H37Rv (NCBI GenBank accession: NC_000962.3). Minority variants were defined as nucleobase positions with an alternate nucleotide, with respect to the reference, with an allelic frequency of 1% to 74%, where the lower range was dependant on the type of polymorphism; small insertion/deletions (indels) and SNPs would each require minimum frequency of more than 1% and more than 3% respectively. Majority variants were defined as nucleobases with an allelic frequency of 75% or more (20). We focused on the drug resistance conferring minority variants in *rpoB, katG, fabG1, inhA, pncA, panD, embB, embA, gyrA, gyrB*, and *rrs* associated with resistance to first-line (isoniazid, rifampicin, pyrazinamide, ethambutol) and key second-line TB drugs (fluoroquinolones and the injectables such as amikacin, capreomycin and kanamycin).

Potential drug resistance conferring minority variants were called using LoFreq v2.1.1 (17, 18) from BAM files produced by Snippy v3.1 (https://github.com/tseemann/snippy). The indel quality score was inserted into the BAM files and potential minority variants (SNPs and indels) were called. We used the strand bias filter to remove any minority variants with a read depth of less than 5 reads, an allelic frequency below 3% and a maximum strain variation over 0.01 (17, 18). Potential minority variants were annotated using SnpEFF v4.3 (21).

### Drug resistance verification and classification

The potential drug resistance conferring minority variants called by LoFreq were manually cross checked with the 2021 WHO Catalogue (22, 23). Only mutations classified as “Associated with resistance” and “Associated with resistance– Interim” in 2021 WHO Catalogue confidence-grading (22, 23) were included in the study. We defined “Associated with resistance” as high confidence resistance associated variants, and “Associated with resistance – Interim” as interim confidence resistance associated variants. The high confidence drug resistance mutations associated minority variants were manually reviewed by direct visualization of the read alignments using Geneious Prime® 2023.0.4.

### Detection of mixed M. tuberculosis strain populations

Mixed population infection was defined as the presence of more than one strain of *M. tuberculosis* during the same disease episode. We detected mixed populations of *M. tuberculosis* in WGS data using QuantTB v1.0 (19) with default settings (>100 SNP differences between strains) and defined the population that was present in the highest proportion as the majority population. If high-confidence drug resistance conferring minority variants were detected in genome sequences of mixed populations, we manually verified the presence or absence of regions of differences (RDs) (24) using Geneious Prime to determine which lineage the minority variants arose from.

### Statistical analysis

Prism GraphPad v9.4.1 was used for statistical analyses. Differences in drug resistant between minority variants and mixed populations were assessed using the Fisher’s exact test with a significance level of p<0.05.

## Results

A total of 1831 *M. tuberculosis* strains sequenced during routine WGS were included in the analysis, which represent the majority of TB cases identified in NSW during the 5-year study period (Figure 1, Supplementary Table S1). In this study, isolates were considered drug resistant if resistance was detected either on routine pDST, or if WHO defined drug resistance associated mutations were identified by WGS (if pDST was absent or isolates were reported susceptible).

**Figure 1:**
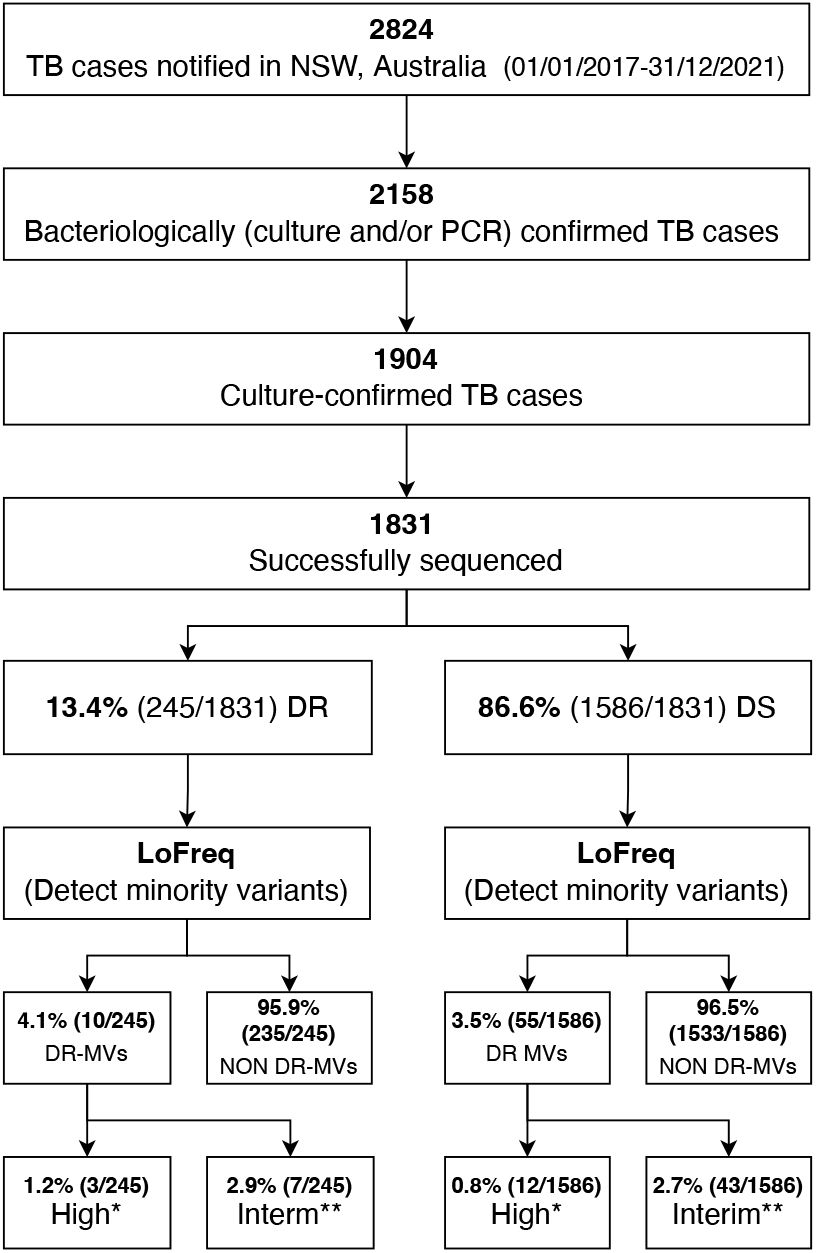
Detection of drug resistance conferring minority variants in all sequenced *M. tuberculosis* strains using LoFreq. DR: drug resistant (majority variants resistant to any first-line TB drugs); DS: drug susceptible (majority variants susceptible to all first-line TB drugs); DR-MVs: drug resistant minority variants; NSW: New South Wales; PCR: *M. tuberculosis*-specific Polymerase Chain Reaction; TB: tuberculosis *High confidence drug resistant minority variants in WHO Catalogue **Interim confidence drug resistant minority variants in WHO Catalogue.

### Detection of drug resistance conferring minority variants

Drug resistance conferring minority variants were detected in 3.5% (65/1831) of sequenced strains (Figure 1); 70.8% were cultured from respiratory specimens (Supplementary Figure S1a). Of the minority variants with drug resistance mutations, 15/65 (23.1%) had high confidence mutations (Table 1) and 50/65 (76.9%) had interim confidence mutations (Supplementary Table S2). All drug resistance associated minority variants identified were supported by at least four overlapping reads with at least one forward and one reverse read; strand bias was less than 0.001. The allelic frequency of minority variants associated with drug resistance varied from 3.3% to 23.0% (Figure 2). Minority variants with drug resistance mutations were found in all major lineages (Supplementary Figure S1b), but more commonly associated with Lineage 1 compared to other lineages, however this was not statistically significant (p=0.08).

**Table 1:** Overview of high confidence drug resistance conferring mutations identified in *M. tuberculosis* minority variants, including mixed strain infections.

| Strain ID | Source | Smear microscopy | Lineage | gDST profile | pDST profile | Mixed strain infection* (Y/N) |
| --- | --- | --- | --- | --- | --- | --- |
| 18-1773-0435 | Pus mediastinal | Negative | Lineage 1 | S | INH-R | Y |
| 17-3391-0281 | Lavage | Negative | Lineage 1 | S | S | Y |
| 17-3391-0008 | Fluid-Neck | Negative | Lineage 1 | S | S | Y |
| 17-3391-0083 | Sputum | Negative | Lineage 2 | fabG1 -15 c>t | INH-R | Y |
| 18-1773-0007 | Sputum | Negative | Lineage 3 | S | S | Y |
| 20-002-0186 | Sputum | Negative | Lineage 1 | fabG1 -15 c>t | INH-R | N |
| 20-002-0197D1 | Sputum | Positive | Lineage 1 | S | S | N |
| 21-002-0305 | Tissue - Pleura | Negative | Lineage 1 | S | S | N |
| 21-002-0237 | Tissue - Lung | Negative | Lineage 1 | S | S | N |
| 17-3391-0013 | Lymph Node | Negative | Lineage 2 | S | S | N |
| 17-3391-0018 | Sputum | Negative | Lineage 2 | S | S | N |
| 21-002-0252D1 | Sputum | Negative | Lineage 2 | S | S | N |
| 18-1773-0451 | Uncertain | Negative | Lineage 4 | S | S | N |
| 20-002-0411 | Washing | Negative | Lineage 4 | S | S | N |
| 21-002-0328 | Fluid - Pleura | Negative | Lineage 4 | S | S | N |

gDST: genotypic drug susceptibility testing; ID: identifier; INH: isoniazid; pDST: phenotypic drug susceptibility testing; R: resistant; S: susceptible to all first-line TB drugs; SNP: Single-nucleotide polymorphisms. \*Mixed strains detected by QuantTB (19).

**Figure 2:**
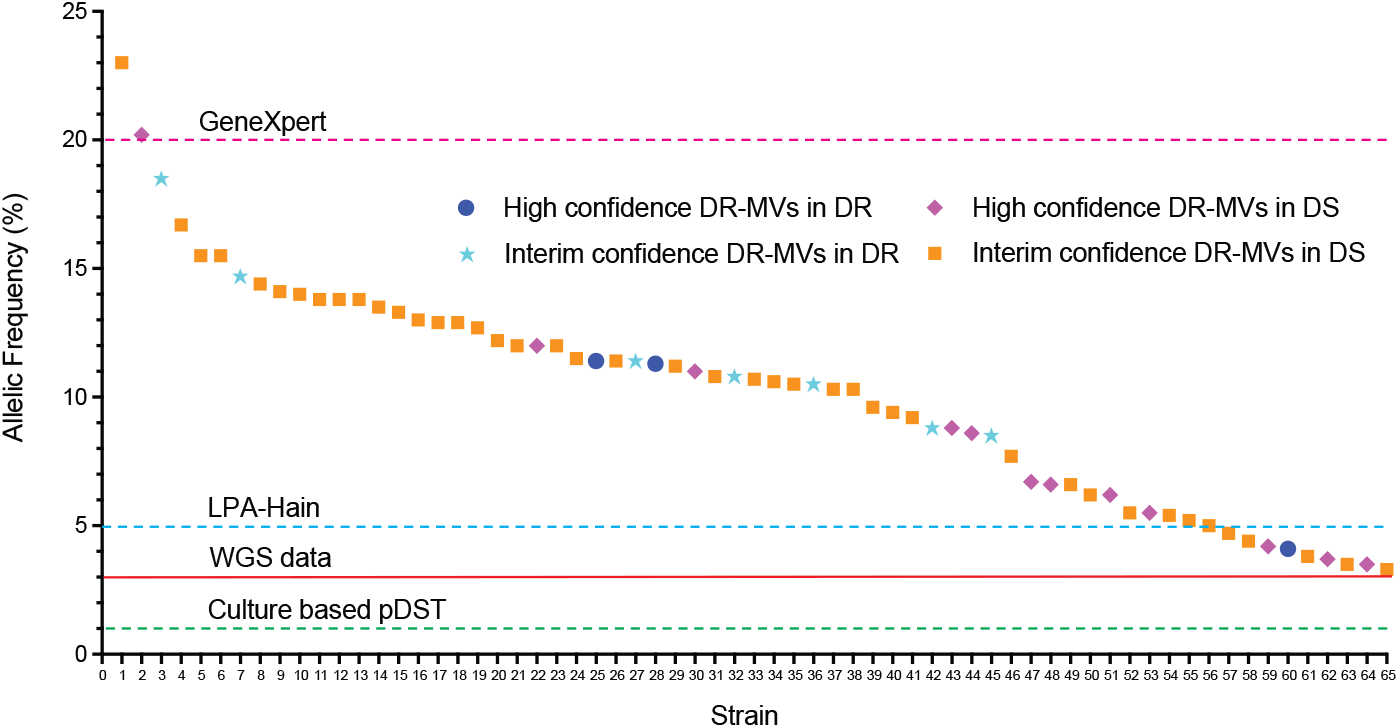
Allelic frequency of drug resistance conferring mutations detected in minority *M. tuberculosis* variants. DR: drug resistant (majority variants resistant to any first-line TB drugs); DS: drug susceptible (majority variants susceptible to all first-line TB drugs); DR-MVs: drug resistant minority variants; LPA: line probe assay; pDST: phenotypic drug-susceptibility testing; WGS: whole genome sequencing. Horizontal red line and dashed lines indicate the limit of detection of minority variants for each method.

The majority of the 65 specimens with drug resistant minority variants (84.6%, 55/65) had majority variants that were phenotypically and genomically drug susceptible. The remaining 10 (15.4%, 10/65) had majority variants with drug resistance along with minority variants with additional resistance (Figure 1, Figure 2). High confidence drug resistance conferring mutations in minority variants were 1.5 times more common when majority variants were drug resistant (1.2%, 3/245) compared to drug susceptible (0.8%, 12/1586; p=0.44).

### Identification of mixed M. tuberculosis strain populations

We identified mixed *M. tuberculosis* strain populations in 183 (10.0%, 183/1831) specimens (Figure 3); 177 specimens were determined to be mixed with two populations (96.7%, 177/183) while six specimens showed signatures of three mixed populations (3.3%, 6/183) (Supplementary Table S1). Specimens with mixed *M. tuberculosis* strain populations were distributed across all four major lineages, but significantly more common in Lineage 3 compared to other lineages (Supplementary Figure S2a) (p=0.0002). The majority of specimens with mixed populations were cultured from respiratory specimens (124/183, 67.8%); 32.2% of non-respiratory specimens had mixed populations (Supplementary Figure S2b).

**Figure 3:**
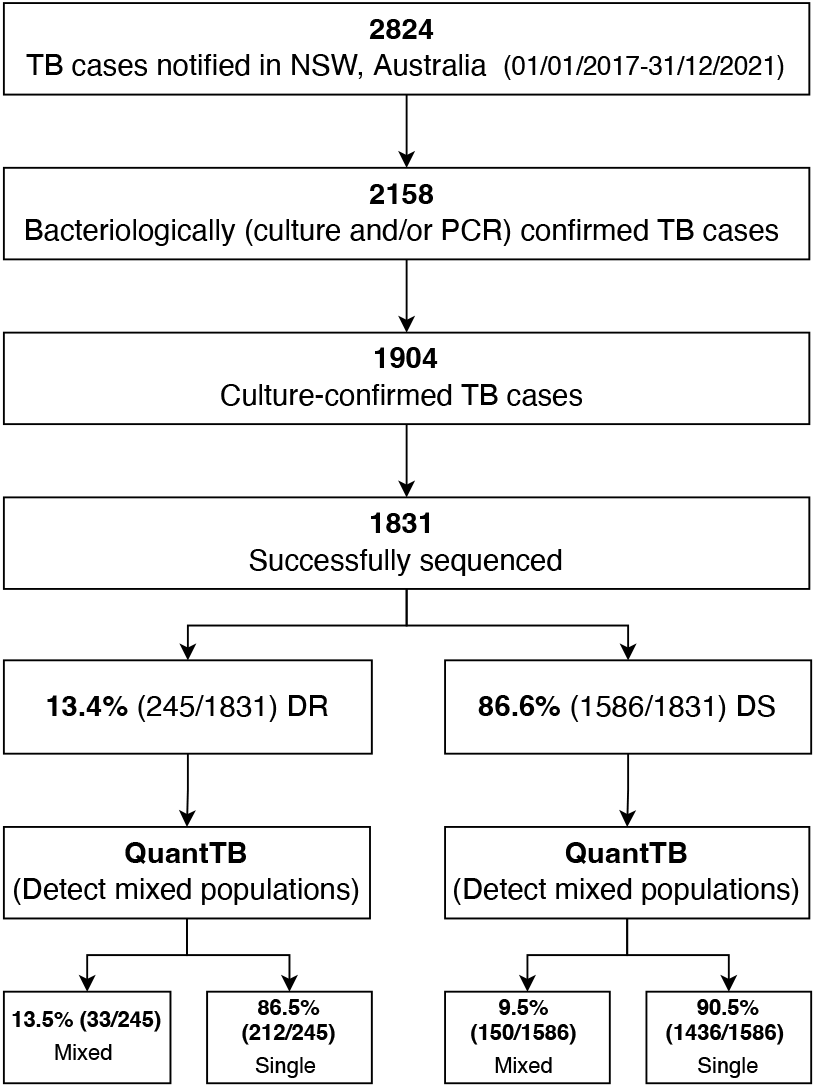
Identification of mixed populations in all sequenced *M. tuberculosis* strains using QuantTB. DR: drug resistant (majority variants resistant to any first-line TB drugs); DS: drug susceptible (majority variants susceptible to all first-line TB drugs); Mixed: mixed populations; NSW: New South Wales; PCR: *M. tuberculosis*-specific Polymerase Chain Reaction; Single: single population; TB: tuberculosis.

Mixed population infections were 42.1% more common in strains where the majority population was drug resistant (13.5%, 33/245) compared to drug susceptible (9.5%, 150/1586; p=0.07). The combined results of LoFreq and QuantTB, to determine the proportion of strains with drug resistant minority variants that represent mixed populations, are represented in Figure 4. Drug resistant minority variants were documented in 6.0% (11/183) of specimens with mixed populations (Figure 4, Figure 5); 0.8% (2/245) where the majority strain was considered drug resistant and 0.6% (9/1586) where the majority strain was considered drug susceptible. While 3.3% (54/1648) of specimens with a single strain population harbored drug resistant minority variants. This occurred in 6.0% (11/183) of those with mixed population infections, regardless of whether the strain was considered drug resistant or susceptible (Figures 4 and 5).

**Figure 4:**
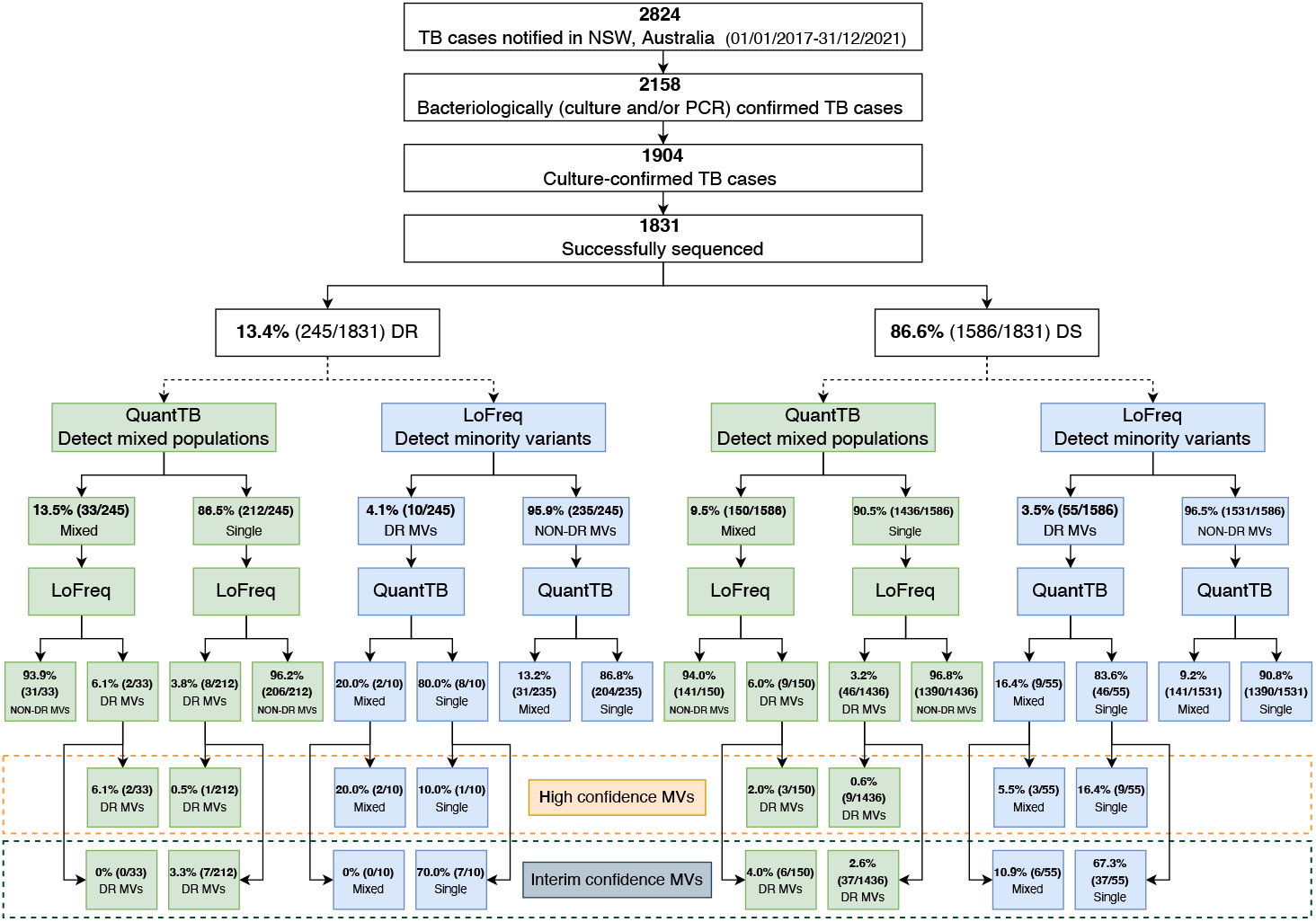
Combined assessment of drug resistant minority variants and mixed *M. tuberculosis* strain populations detected in specimens with majority drug resistant or drug susceptible strains. DR: drug resistant (majority variants resistant to any first-line TB drugs); DS: drug susceptible (majority variants susceptible to all first-line TB drugs); DR-MVs: drug resistant minority variants; MDR: multidrug resistant; Mixed: mixed populations; NSW: New South Wales; PCR: *M. tuberculosis*-specific Polymerase Chain Reaction; Single: single population; TB: tuberculosis.

**Figure 5:**
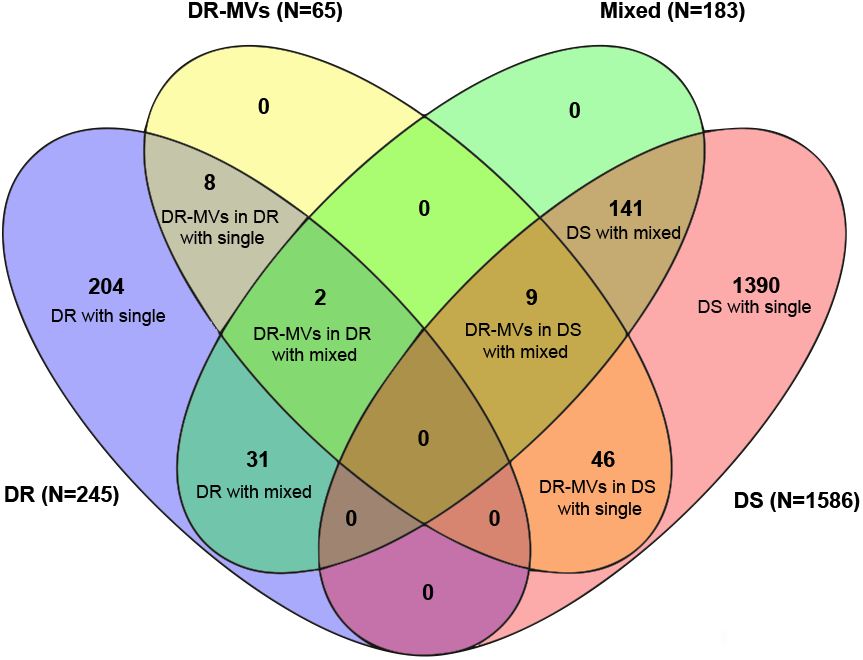
Overview of *M. tuberculosis* drug resistant minority variants and mixed strain populations detected. DR: drug resistant (majority variants resistant to any first-line TB drugs); DS: drug susceptible (majority variants susceptible to all first-line TB drugs); DR-MVs: drug resistant minority variants; Mixed: mixed populations; Single: single population. Numbers (N) in brackets next to legend indicate the total number of isolates per legend.

### Assignment of drug resistance conferring minority variants

The assignment of high confidence drug resistance mutations detected in minority variants in single strain population specimens are summarized in Table 2 (interim confidence mutations in Supplementary Table S2). One specimen considered phenotypically and genotypically isoniazid mono-resistant (majority variant *fabG1* -15 c>t) also had a *rpoB*_H445N mutation detected as a minority variant at 11.4% allelic frequency, indicating potential for selection of this MDR strain with inappropriate treatment (Table 2). Minority variants with high-confidence drug resistance conferring mutations were more frequently detected in specimens with mixed strain populations (2.7%, 5/183) than in those with a single strain population (0.6%, 10/1648) (p=0.01). Due to limitations of the bioinformatics software and short-read sequencing technology used, minority drug resistance mutations were unable to be assigned to either the minority or majority population within specimens with mixed population (Table 3).

**Table 2:**
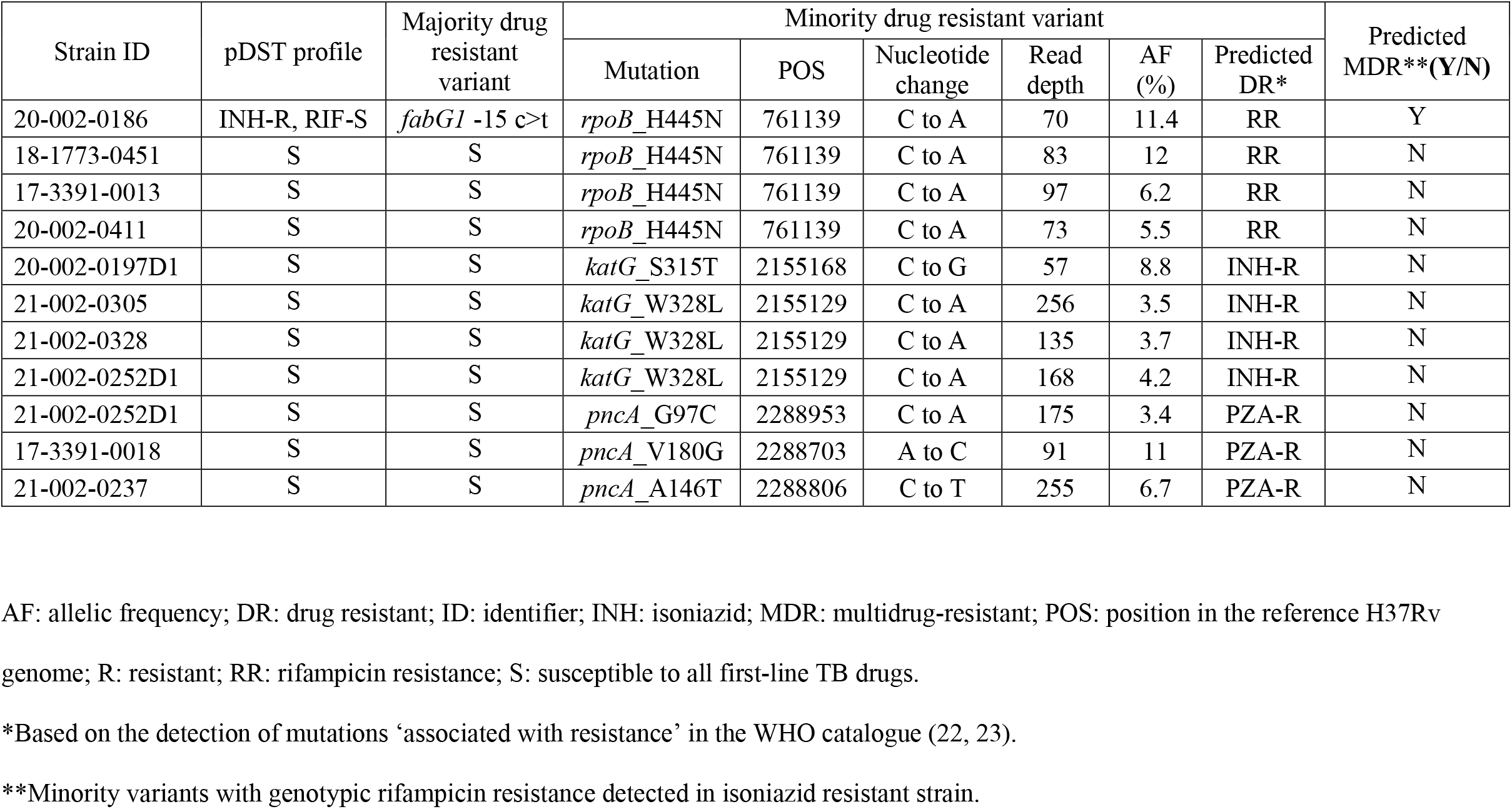
High confidence drug resistance conferring mutations identified in minority variants from specimens with a single *M. tuberculosis* strain (N=1648)

**Table 3:** High confidence drug resistance conferring mutations identified in minority variants from specimens with more than one *M. tuberculosis* strain (N=183)

| Strain ID | pDST profile | Majority drug resistant variant | Mixed populations ratio | Minority drug resistant variant |  |  |  |  |  |
| --- | --- | --- | --- | --- | --- | --- | --- | --- | --- |
|  |  |  |  | Mutation | POS | Nucleotide change | Read depth | AF (%) | Predicted DR** |
| 17-3391-0083 | INH-R, RIF-S | *fabG1* -15 c>t | 63% : 37% | *rpoB*_H445N | 761139 | C to A | 222 | 4.1 | NA |
| 18-1773-0435 | INH-R | S* | 81.9% : 18.1% | *katG*_S315T | 2155168 | C to G | 62 | 11.3 | NA |
| 18-1773-0007 | S | S | 80.4% : 19.6% | *katG*_S315T | 2155168 | C to G | 89 | 20.2 | NA |
| 17-3391-0281 | S | S | 57.9% : 42.1% | *pncA*_V128G | 2288859 | A to C | 244 | 6.6 | NA |
| 17-3391-0008 | S | S | 51.7% : 48.3% | *embB*_D354A | 4247574 | A to C | 140 | 8.6 | NA |

AF: allelic frequency; DR: drug resistant; EMB: ethambutol; ID: identifier; INH: isoniazid; L: lineage; NA: not applicable; POS: position in the reference H37Rv genome; PZA: pyrazinamide; R: resistant; RR: rifampicin resistance; S: susceptible to all first-line TB drugs.
\*No isoniazid associated resistance mutation detected.
\*\*Minority variants were not able to be assigned to either the majority population or minority population within the isolates.

## Discussion

This study is the first to quantify the frequency with which minority variants that harbour mutations conferring drug resistance are observed in routinely sequenced isolates from a low TB incidence setting. It is also the first to differentiate same strain minority variants from those representing mixed *M. tuberculosis* strain populations. From a programmatic perspective, the assessment of drug resistant minority variants improves clinical risk management and should provide patient benefits, especially in settings where disease relapse is a concern.

Drug resistance conferring minority variants are not routinely interrogated during WGS-based gDST, but may be detected if pDST is performed, leading to discrepancies between WGS and phenotypic data. Our analysis of WGS data from culture-confirmed clinical *M. tuberculosis* strains confirmed previous findings that minority strains carrying high confidence drug resistance mutations (such as *katG*_S315T at 11.3% allelic frequency) can account for occasional discrepancies between pDST and gDST (7, 25). Routinely assessing for minority variants with drug resistance conferring mutations in clinical *M. tuberculosis* specimens would enhance drug resistance surveillance and have particular clinical relevance to prevent drug resistance amplification and disease relapse.

Minority variants with drug resistance conferring mutations, especially in strains demonstrating phenotypic drug resistance, may potentially develop additional drug resistance if exposed to sub-optimal treatment. Therefore, the presence of these minority variants provides an early risk indication for treatment failure (6, 7, 16, 26). The presence of minority variants with additional resistance to rifampicin or isoniazid are particularly important when majority variants have mono-resistance to either of these first-line drugs, as these strains would then become multidrug resistant (MDR). In our analysis, we identified the presence of minority variants with additional rifampicin resistance in 1/243 (0.4%) of TB cases with isoniazid monoresistance. However, genomically and phenotypically undetectable MDR minority variants may be more common in high incidence settings. The early detection of drug resistant minority variants should guide optimal treatment to reduce the risk of failure or relapse.

Our findings confirmed previous observations that minority *M. tuberculosis* variants with drug resistance conferring mutations can be detected by WGS, even when this is not apparent on pDST (4). Even if not MDR, unrecognised minority variants with drug resistance could be selected during sub-optimal treatment, leading to treatment failure and clinical relapse (27). The sensitivity of pDST has been reported to be as high as 1% of the strain population (8, 10), however, our gDST data reflected instances where 6.6% of strains had high confidence drug resistance mutations but still tested susceptible on pDST. In total 3.5% (55/1586) of strains that tested drug susceptible on pDST had minority variants with drug resistance associated mutations. These discrepant results warrant further interrogation as undetected drug resistance have important laboratory reporting and clinical management implications.

Some limitations of our analysis need to be acknowledged. Laboratory subcultures of *M. tuberculosis* can reduce clonal diversity, with culture selection leading to a loss of the information on resistant minority populations in clinical samples (28, 29). The drug resistant minority variants observed from culture-derived WGS data, may not capture all relevant *M. tuberculosis* population dynamics (6). Culture independent sequencing, without any pre-selection from culture, may be more reflective of the bacterial population in the clinical sample.

We synergistically employed two tools, LoFreq to determine minority variants associated with drug resistance and QuantTB to detect mixed *M. tuberculosis* strain infections. However, neither software was able to assign minority variants associated with drug resistance to majority or minority populations within the isolate. It is possible that the drug resistant minority variant belongs to the minority population, due to the fact that the vast majority of the strains tested were phenotypically and genotypically consistent. However, it is also possible that the minority drug resistant variants arose from the majority strain population, but were not detected using pDST. For instance, if the minority variant, *rpoB*_H445N, was present in the majority population of our sample with a majority variant *fabG1* C-15T, there would be the potential risk of developing MDR-TB. In addition to the limitations with bioinformatic pipelines, the use of the short-read sequencing technology can also impact the assignment. It is unlikely to capture both drug resistant minority variants and strain population differences in a single 150 bp read. These highlights the need for bioinformatic pipelines or long-read sequencing technology with the ability to assign drug resistance conferring minority variants to either the majority or minority populations within the genome sequence of *M. tuberculosis* from a clinical sample.

The allelic frequency of minority variants with resistance detected in our analysis ranged from 3.3% to 23.0%. Previous studies suggested that the presence of drug resistant minority is associated with poor treatment outcomes (6, 30), especially if these minority variants have an allelic frequency of more than 5% (16, 30). However, some studies indicate that even at very low (<1%) allelic frequencies may be increased during sub-optimal treatment leading to fixed resistance and a gain in fitness (14, 31).

In conclusion, our findings demonstrate that minority variants of *M. tuberculosis* with drug resistance conferring mutations can be detected using routine WGS. Minority variants with high-confidence drug resistance conferring mutations were more prevalent in patients with mixed strain populations. The high-resolution interrogation of drug resistant minority variants could improve clinical risk assessment and strengthen surveillance of drug resistant strains.

## Data availability

Raw de-identified pathogen WGS data was deposited in the National Center for Biotechnology Information (NCBI) Short Read Archive (SRA) under BioProject number PRJNA899911.

## Acknowledgements

The authors thank the Sydney Informatics Hub and the University of Sydney’s high-performance computing cluster, Artemis. The first author (X.Z.) is funded by NHMRC Centre for Research Excellence in Tuberculosis. The funders of this study had no role in the study design, data collection, data analysis and interpretation, or writing of the report. The corresponding author had full access to study data and final responsibility for the decision to submit for publication.

## Author contributions

B.J.M. and V.S. designed the study and guided the data analysis. Data collection was done by T.C., E.M., C.L., and E.S. collected the data. X.Z. performed bioinformatic analysis and wrote the first manuscript draft with editing from all authors. The final manuscript was approved by all authors.

## Supplemental Material

**Figure S1:**
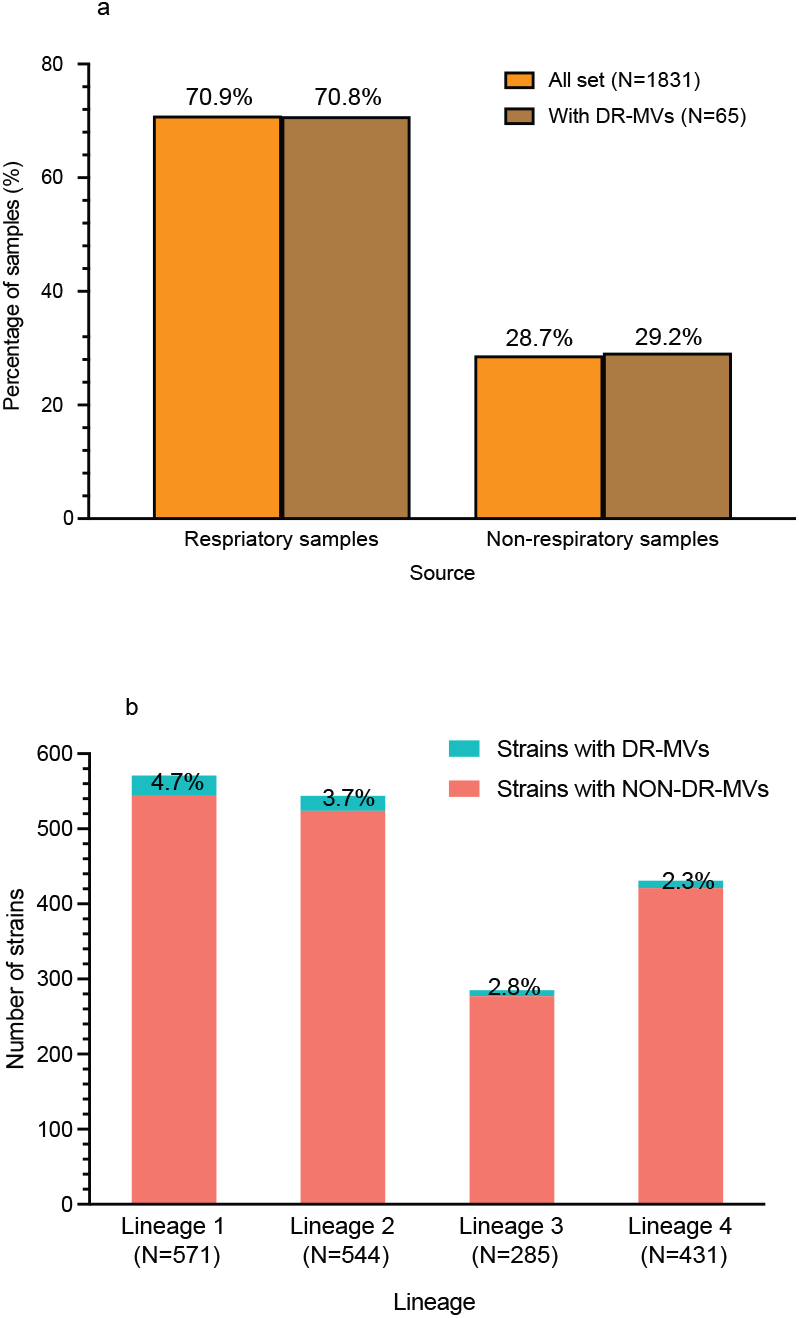
*M. tuberculosis* strains with drug resistance conferring minority variants; (a) the percentage of respiratory and non-respiratory specimens, (b) the number and percentage of strains with minority variants detected in all four major lineages. DR-MVs: drug resistant minority variants. Numbers (N) in brackets next to lineages (X axis) indicate the total number of isolates per lineage.

**Figure S2:**
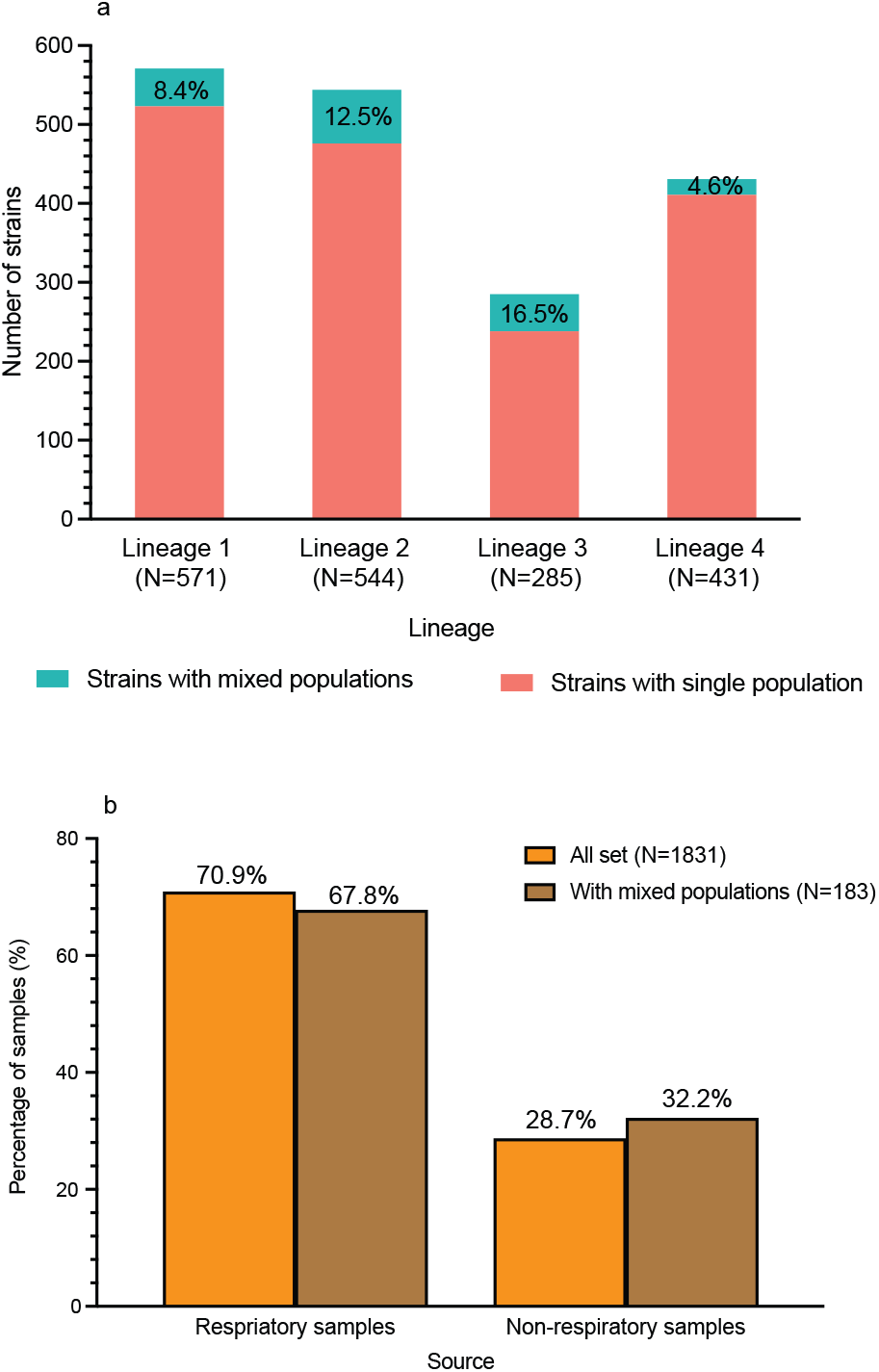
*M. tuberculosis* strains with mixed populations; (a) the number and percentage of mixed strain populations detected in different *M. tuberculosis* lineages, (b) the percentage in respiratory and non-respiratory specimen sources. Numbers (N) in brackets next to lineages (X axis) indicate the total number of isolates per lineage.

